# A novel rRNA hybridization-based approach to rapid, accurate *Candida* identification directly from blood culture

**DOI:** 10.1101/2022.05.09.491195

**Authors:** Michelle E. Matzko, Poppy C. S. Sephton-Clark, Eleanor L. Young, Tulip A. Jhaveri, Melanie A. Martinsen, Evan Mojica, Rich Boykin, Virginia M. Pierce, Christina A. Cuomo, Roby P. Bhattacharyya

## Abstract

Invasive fungal infections are increasingly common and carry high morbidity and mortality, yet fungal diagnostics lag behind bacterial diagnostics in rapidly identifying the causal pathogen. We previously devised a fluorescent hybridization-based assay to identify bacteria within hours directly from blood culture bottles without subculture, called phylogeny-informed rRNA-based strain identification (Phirst-ID). Here, we adapt this approach to unambiguously identify 11 common pathogenic *Candida* species, including *C. auris*, with 100% accuracy from laboratory culture (33 of 33 strains in a reference panel, plus 33 of 33 additional isolates tested in a validation panel). In a pilot study on 62 consecutive positive clinical blood cultures from two hospitals that showed yeast on Gram stain, *Candida* Phirst-ID matched the clinical laboratory result for 58 of 59 specimens represented in the 11-species reference panel, without misclassifying the 3 off-panel species. It also detected mixed *Candida* species in 2 of these 62 specimens, including the one discordant classification, that were not identified by standard clinical microbiology workflows; in each case the presence of both species was validated by both clinical and experimental data. Finally, in three specimens that grew both bacteria and yeast, we paired our prior bacterial probeset with this new *Candida* probeset to detect both pathogen types using Phirst-ID. This simple, robust assay can provide accurate *Candida* identification within hours directly from blood culture bottles, and the conceptual approach holds promise for pan-microbial identification in a single workflow.

## Introduction

Invasive fungal infections (IFIs) are a leading cause of morbidity and mortality, accounting for at least 1.6 million global deaths annually (1, 2) and rising in incidence and severity in recent years (2–4), particularly in growing immunocompromised populations. Despite this critical threat, clinical fungal diagnostics remain slow and cumbersome (5–8), leading to delayed recognition and treatment with clear mortality cost (7, 9, 10) and increased empiric antifungal use (11). Rapid assays for fungal biomarkers such as the cell wall components beta-D-glucan or galactomannan have emerged in recent years, but these largely lack the sensitivity or specificity to guide clinical care, aside from antigen tests for a limited number of species (5, 6, 8). Traditional methods like histology and culture thus remain the gold standard for clinical microbiology laboratories despite a number of drawbacks, including low sensitivity, slow turnaround time, and the need for considerable laboratory infrastructure and expertise (5, 6, 8, 12).

One critical subset of IFIs, *Candida* bloodstream infections, exhibit high mortality (13–15) and rising incidence due in part to growth of immunosuppressed populations over time (2, 3). Despite recent advances in candidemia diagnostics, and one FDA-cleared assay to detect *Candida* species in uncultured blood (16), accurate species identification in candidemia remains slower than in bacteremia in most clinical settings, due in part to slower growth on the subculture step required prior to mass spectrometry-based identification (17).

We recently developed a novel approach to rapid, sensitive bacterial identification directly from primary clinical samples or cultured blood, based on a robust, commercially available assay platform (NanoString) that enables multiplexed hybridization to highly abundant ribosomal RNA (rRNA) (18). This assay, which we termed phylogeny-informed rRNA-based strain identification (Phirst-ID), effectively decodes sequence differences of highly conserved 16S and 23S rRNA subunits between species based on differential hybridization to a fixed set of probes. Because it relies only on rRNA sequence, the physiological state of the organism does not affect results, meaning that it can work directly on clinical samples without subculture. The abundance of the rRNA targets also enhances sensitivity: we showed that the assay platform can detect as few as 10 bacteria in a sample (19), facilitating implementation on primary clinical samples. In principle, this assay should extend to any pathogen with rRNA sequences unique enough to distinguish individual species.

Here, we extend this Phirst-ID approach to fungal pathogens by demonstrating that we can sensitively identify major clinically relevant *Candida* species. We designed a 21-probe set targeting the 18S and 28S rRNA subunits that accurately identifies 11 *Candida* species, including *C. auris*, with 100% accuracy at the species level on a reference panel of 33 isolates grown in axenic culture. After validating the *Candida* Phirst-ID assay on this reference panel and an independent validation set of 33 isolates grown in laboratory culture, we assessed its performance on 62 consecutive positive blood cultures from two collaborating hospitals that showed yeast forms on Gram stain. Phirst-ID matched the species identified by the standard clinical microbiology workflow for 58 of 59 specimens harboring *Candida* species from our reference panel, returning results in less than 8 hours, with less than 30 minutes of hands-on time. The one discordant classification was a mixed culture of *C. albicans* and *C. dubliniensis*; only the *C. albicans* was found by the clinical microbiology laboratory, whereas Phirst-ID identified *C. dubliniensis*.

## Materials and Methods

### Probeset design

We collated the full 18S and partial 28S rRNA sequences for 11 medically relevant *Candida* species *(albicans*, *auris*, *dubliniensis*, *duobushaemulonii*, *glabrata*, *guilliermondii*, *haemulonii*, *krusei*, *lusitaniae*, *parapsilosis*, and *tropicalis*) using publicly available genomes (**Supplementary Table S1**). We also compiled 18S rRNA sequences from 112 medically relevant or environmental non-*Candida* fungi from the Silva database (20) to serve as an outgroup for probe design, to avoid unintentional cross-reactivity against other fungi that might be present in clinical samples. Since 28S rRNA data are less well annotated across non-pathogenic fungi, we did not include them among our outgroup sequences. We then profiled each desired target sequence using NanoString’s probe design algorithm to identify all putative pairs of consecutive 50mer probe-binding regions within the 18S and 28S subunits of each species that meet parameter specifications for predicted binding kinetics and thermodynamics, secondary structure, and sequence composition for the desired targets while minimizing predicted cross-reactivity against non-targeted species within these databases, or the human genome (18, 21). The resulting probeset is shown in **Supplementary Figure S1**; species-specific probes tended to cluster in variable regions of each rRNA subunit and often overlapped. Of the 11 desired species targeted by this approach, we successfully designed probes to 10 of the 11 18S rRNA regions and 9 of the 11 28S rRNA regions; for *C. guilliermondii*, only an 18S probe was predicted to be sufficiently species-specific, and for *C. dubliniensis*, no species-specific probes could be designed to either target due to its high similarity to *C. albicans*, *C. parapsilosis*, and *C. tropicalis* in both subunits. We did not target internal transcribed spacer (ITS) regions, despite their greater inter-species variability, due to their lower steady-state abundance, which would reduce the sensitivity of detection (22). We also designed probes to each rRNA subunit intended to recognize all members of the *Candida* genus while excluding other genera among the outgroups we considered. Probe sequences are listed in **Supplementary Table S2**.

### Strain acquisition

The majority of the 66 *Candida* isolates used in this manuscript were obtained from the CDC Antibiotic Resistance (AR) Isolate Bank (23). When triplicate strains of the same species were not available from the CDC AR Bank, they were supplemented from strains on hand at the Brigham & Women’s Hospital (BWH) or Massachusetts General Hospital (MGH) Clinical Microbiology laboratories (**Supplementary Table S3**).

### Clinical sample processing

We collected 1 mL of broth from 62 consecutive blood culture bottles that signaled positive and showed yeast on Gram stain in the clinical microbiology laboratory at MGH or BWH between September 2020 and June 2021. Samples were set aside at 4°C in the clinical microbiology laboratory until they were picked up by research staff, typically within 24 hours of signaling positive, then stored at −80°C before being processed in batch. Batches of 12 samples of blood culture broth were thawed on ice, then lysed, detected, and analyzed as described below. Collection of these discarded clinical samples was approved under waiver of consent by the Mass General Brigham Institutional Review Board (IRB), which governs both hospitals, under protocol number 2015P002215. No results were returned to clinicians, but species identifications generated by Phirst-ID were compared to the results of standard workflows in the clinical microbiology laboratories, as approved by the IRB. Researchers were blinded to all clinical results until a Phirst-ID species identification was made.

### Lysate preparation

Frozen strains were freshly streaked on yeast extract-peptone-dextrose (YPD) agar plates at 30°C for ∼72 hours until colonies were ∼2-3mm in size, then a portion of one colony was incubated in 4mL of Roswell Park Memorial Institute (RPMI) media for ∼12-24 hours at 30°C to logarithmic phase (optical density at 600 nm of 0.3-0.8). Fresh blood culture broth from clinical samples was diluted 1:10 in phosphate-buffered saline (PBS). 200uL of either sample type was mixed with 200uL of RLT buffer (Qiagen) with 1% ß-mercaptoethanol (ß-ME), an irreversible chemical RNase inhibitor, then lysed via mechanical disruption on a FastPrep bead-beater (MP Biosciences) using 200uL of 0.1um zirconia-silica beads (BioSpec) for two 90-second cycles at 10m/s separated by 30 seconds on ice. Lysates from laboratory cultures were diluted 1:10 prior to NanoString; lysates from clinical blood cultures were run as-is.

### NanoString data generation

Lysates were run with our custom-designed *Candida* rRNA probeset (CandID4) on the NanoString Sprint platform using a modified assay protocol as described (18). Briefly, lysates were heated to 95°C for 2 minutes immediately prior to hybridization for 1 hour to facilitate denaturation of rRNA secondary structure and dissociation from ribosomal proteins, accelerating hybridization from the overnight incubation in the standard protocol. Raw binding data (counts per probe) were extracted using NanoString nSolver software (v4.0), and normalized for lane-to-lane variation using positive and negative control spike-ins supplied by NanoString using custom scripts in R (v3.6.1) (https://github.com/broadinstitute/FungalDx_CandidaID). Seven blank samples of RPMI, handled identically to cultured samples, were run to determine background signal for each probe, and were averaged and subtracted from each unknown sample. These normalized, blank-subtracted counts were used for all subsequent analyses.

### Species classification/analysis

Three isolates of each of the 11 *Candida* species to be identified were selected to generate a reference dataset of probeset reactivity profiles (PSRP) for all probes against each strain. Binding patterns for this reference panel were compared to the binding patterns of samples to be identified. To identify species, we used normalized blank-subtracted probe counts to generate heatmaps and pairwise Pearson correlation coefficients using custom scripts in R (https://github.com/broadinstitute/FungalDx_CandidaID). To compute Pearson correlation coefficients, only the 19 species-specific probes were used, since the two pan-*Candida* probes had similar reactivity against all species. The reference panel species with the highest correlation coefficient against a given sample was taken as the predicted species of the comparator sample (18). These predictions were then compared with the species identified through conventional means, either from a reference strain collection or through standard workflows in the clinical microbiology laboratory at the hospital from which the sample was obtained, and classified as either a species match, or a mismatch. Confidence intervals for concordance assessment were calculated using Jeffreys method (24).

### Limit of detection/distinction

To estimate the limit of detection, we performed serial 1:10 dilutions of each species in PBS. At each dilution, we ran Phirst-ID and plated an aliquot (or an appropriate dilution thereof) for colony forming units (cfu) to ascertain the number of cell equivalents in the assay. For each tested species, we estimated the limit of detection as the cfu per assay of the lowest dilution at which probe counts from the most abundant species-matched probe exceeded the average signal of the same probe from mock lysates made from blank RMPI, plus three standard deviations. In addition, we assessed the lowest cfu per assay at which each *Candida* species could be uniquely identified by its Pearson correlation coefficient with our reference panel (limit of distinction).

### Mixed clinical blood cultures

For clinical samples that appeared from manual inspection to have binding signals that suggested the presence of multiple yeast species, an aliquot from the blood culture broth was streaked on ChromAgar *Candida* plates (Becton Dickinson) and inspected for distinct colony morphology and color. Distinct-appearing colonies were restreaked on ChromAgar *Candida* plates until each plate showed uniform colony morphology that matched the original colony, then individual colonies from these uniform plates were grown in liquid RPMI medium and processed and analyzed as above. For specimens from which both fungal and bacterial organisms were identified in the clinical microbiology laboratory, as well as controls without bacterial growth, we separately ran aliquots of blood culture broth on the NanoString using our 21-plex *Candida* Phirst-ID probeset, and the 180-plex bacterial Phirst-ID probeset we previously validated (18). Bacterial probeset reactivity profiles were compared with our previously published reference dataset (18).

### Sanger sequencing

To clarify Phirst-ID findings for three blood culture specimens (BFx121, BFx143, and BFx149), we sequenced two regions of the ITS (25) from representative isolates on ChromAgar *Candida* plates by growing individual colonies in liquid culture, bead-beating, and extracting gDNA using theQuick-DNA/RNA™ Miniprep Plus Kit (Zymo Research). Samples were submitted for Sanger sequencing (GeneWiz) and the closest match was found to a subset of the SILVA database (20) using the BLAST algorithm (26).

## Results

### Common *Candida* species are distinguished by unique probeset reactivity profiles

We compiled publicly available sequence data for the 18S and 28S rRNA subunits from 11 medically relevant *Candida* species, and from 112 non-*Candida* fungi as outgroups to avoid cross-reactivity (**Supplementary Table 1**). We designed 19 probes uniquely targeting species-specific variable regions of the 18S and 28S rRNA subunits from 10 of the medically relevant *Candida* species (**Supplementary Figure S1**). We were unable to design probes predicted to specifically recognize the *C. guilliermondii* 28S subunit, or either subunit for *C. dubliniensis*, due to their similarity with other species (see Methods for details). However, we included these species in subsequent experiments, since our experience with bacterial species suggested that the overall probeset reactivity profile (PSRP) may still uniquely identify closely related species, even without a species-specific probe (18). In addition, we designed two genus-level “pan-*Candida*” probes intended to recognize the 18S and 28S subunits from all *Candida* species while excluding non-*Candida* fungi.

We first tested a reference panel of 3 isolates from each of 11 medically relevant *Candida* species (**Figure 1**; **Supplementary Table S3**). All species were recognized by the two pan-*Candida* probes; aside from those, the majority of species were most strongly recognized by the species-specific probes that were designed to target their species (**Figure 1a**). As we saw with bacterial probes, some probes designed to be species-specific showed some degree of cross-reactivity with phylogenetically similar species (e.g., *C. parapsilosis* and *C. tropicalis*). However, in aggregate, the full PSRP was distinct for each species. Phylogenetically similar strains that are critical to diagnose at the species level but can confound standard microbiological assays, such as *C. auris* versus *C. haemulonii* or *C. duobushaemulonii*, were readily distinguishable.

**Figure 1.**
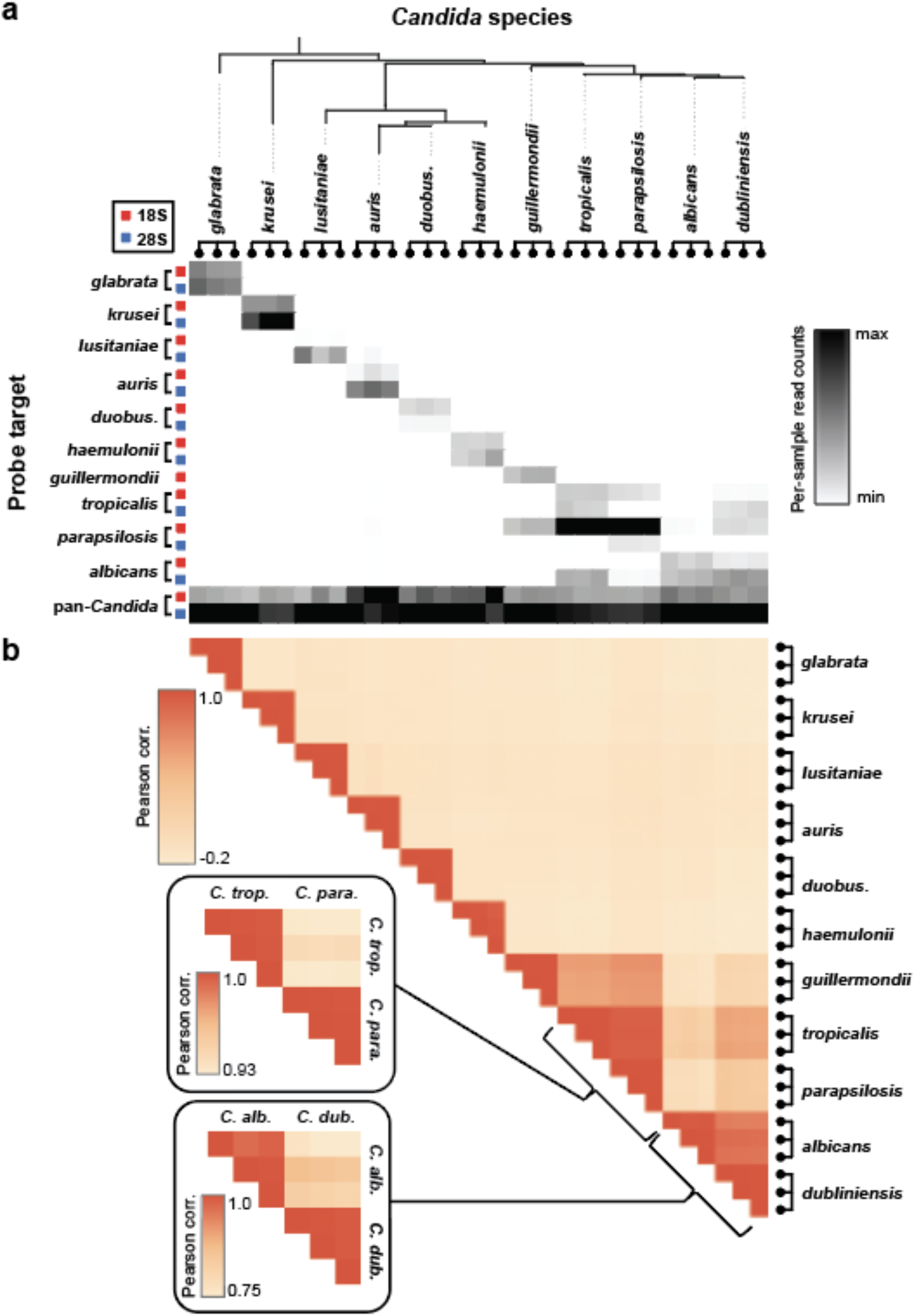
*Candida* Phirst-ID panel uniquely recognizes each of 11 *Candida* species. **(a)** Heatmap of normalized, background-subtracted binding intensities of 21 probes designed to target either the 18S or 28S rRNA subunit from three independent isolates of each of 11 *Candida* species, ordered by phylogenetic similarity. **(b)** Heatmap of Pearson correlation coefficients of the probeset reactivity profiles from 19 species-specific probes. Insets are rescaled for the indicated subset of strains.

### Pearson correlations accurately distinguish each species

To quantify differences in the overall PSRP by species, we calculated a pairwise Pearson correlation coefficient for each sample with each other strain tested in the reference set, as we did for bacterial Phirst-ID (18). This simple analytical approach incorporates data from all probes tested and matches the entire reactivity profile; the strongest correlations should exist within species, whereas non-species-matched pairs should have lower correlation coefficients. We then performed a leave-one-out analysis on this reference panel, comparing the Pearson correlation of each isolate with all other PSRPs in the reference set (aside from its own). In each case, the two closest matches for each isolate in the reference panel were with the other two isolates of the same species (**Figure 1b**; **Supplementary Figure 2**). For example, *C. tropicalis* and *C. parapsilosis* have visually similar PSRPs, yet their Pearson correlation coefficients vary enough that they are clearly distinguishable. In all, 33 of 33 isolates were correctly identified in this manner (100% accuracy; 95% confidence interval 93-100%).

As validation, we tested an additional 33 *Candida* isolates from 7 species by growing them in laboratory culture, lysing, detecting on NanoString (**Supplementary Figure S3**), and comparing them to our reference panel using Pearson correlations (**Figure 2**). In each case, the correct species was identified by the member of the reference panel with the highest correlation (100% accuracy; 95% confidence interval 93-100%) - in fact, the top 3 matches in all 33 cases were to the correctly matched species.

**Figure 2.**
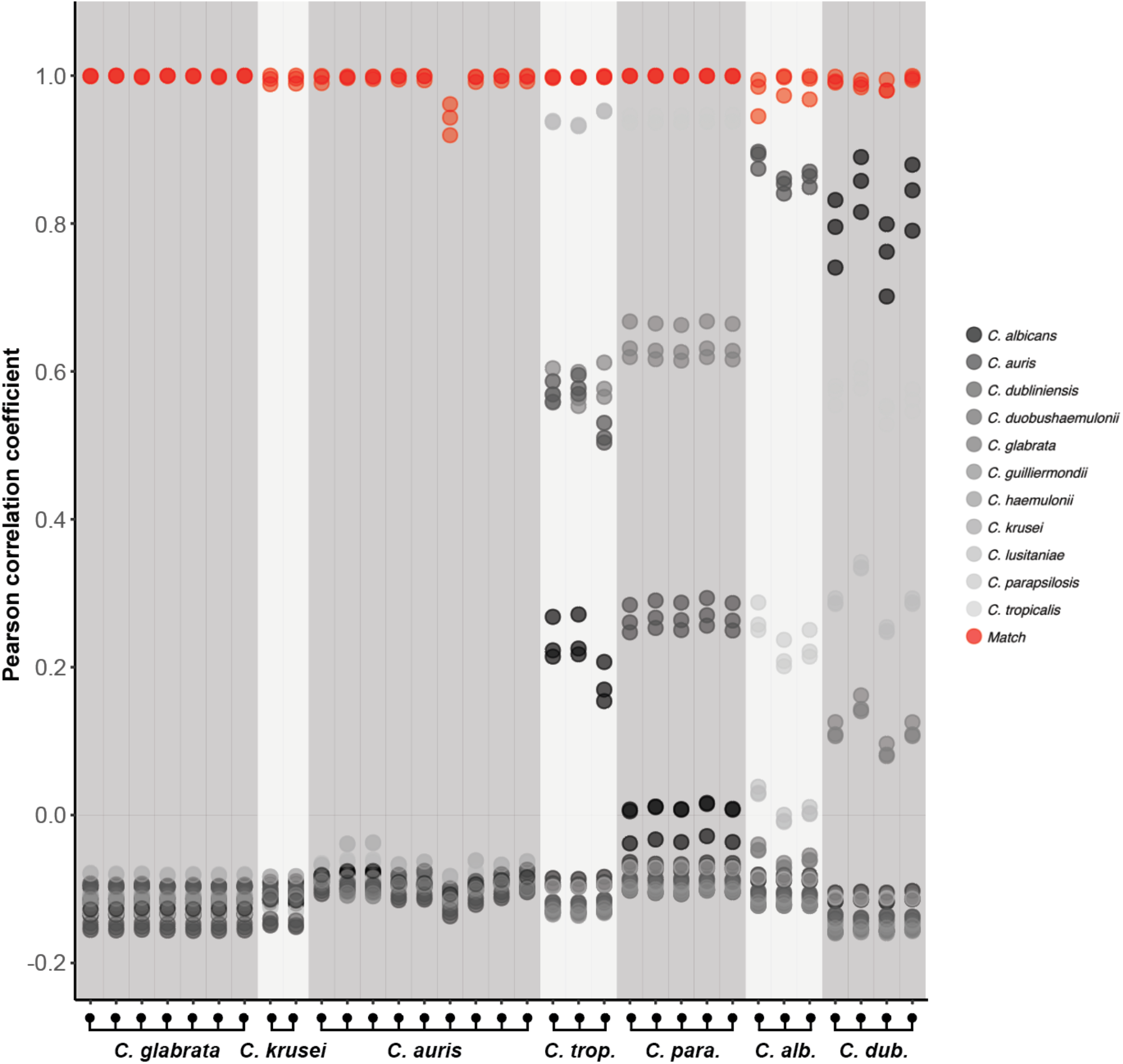
Phirst-ID probeset reactivity profile identifies *Candida* species based on similarity to the reference panel. Pearson correlation coefficients computed from Phirst-ID of 33 independent *Candida* isolates from 7 different species compared with each of 33 *Candida* isolates in a reference panel. Shading of data points indicate the comparison species from the reference panel; species matches are shown in red.

### Assay limit of detection averages around a single yeast cell

To assess limit of detection (LOD), we grew *C. albicans*, *C. auris*, *C. glabrata*, *C. parapsilosis*, and *C. tropicalis* in RPMI media, then performed 1:10 serial dilutions in PBS down to 1:100,000, spanning 5 orders of magnitude. We confirmed the number of yeast cells present at each dilution for each species by plating and counting colony forming units (cfu). We defined the LOD of our assay as the cfu count at which the mean signal for the species-specific probe exceeded that from 7 runs of RPMI controls (mean plus three standard deviations). By this method, we determined our LOD to be ∼1 cfu per assay for most species (**Figure 3**, filled circles). Since lysis occurred at >100x the volume that was run on each NanoString assay, errors related to Poisson statistics at very low cell numbers were minimized, and the LOD could theoretically be less than one cell equivalent per assay. The dilutions and LOD were similar both for replicates of the same species, and across all five *Candida* species tested. While some *Candida* species could be detected at <1 cfu per assay, the number of cfu required to correctly identify each species by Pearson correlation, which we term the “limit of distinction”, was ∼10-fold higher, ranging from 1.5 to 29 cfu (**Figure 3**, red circles). Limits of detection and distinction for each species are tabulated in **Figure 3f**.

**Figure 3.**
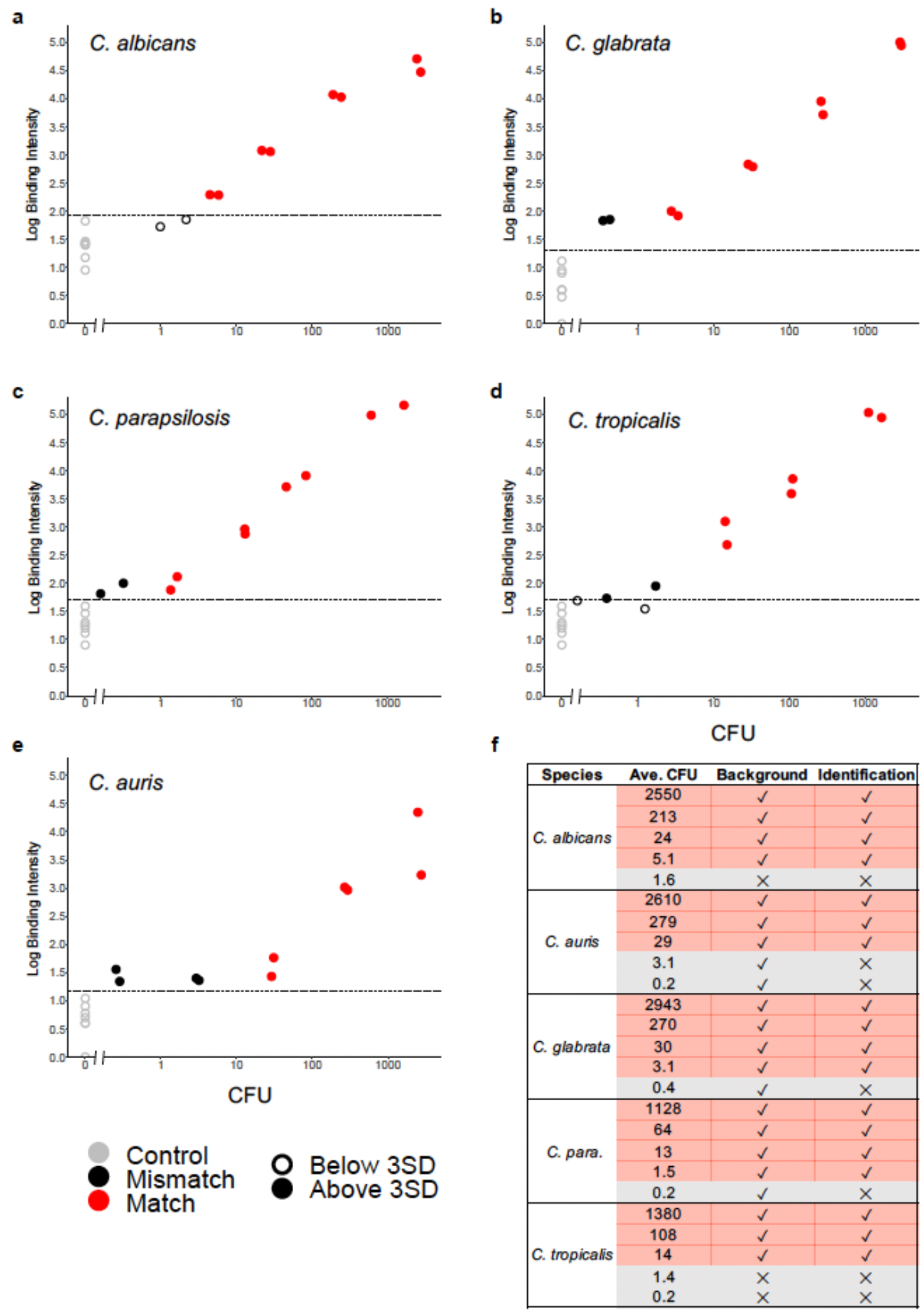
Phirst-ID limits of detection and distinction for 5 *Candida* species. **(a-e)** Serial dilutions of the indicated *Candida* species were tested by Phirst-ID and plated for cfu (x-axis). Normalized binding intensity (y-axis) of the strongest species-specific probe for each sample was compared with that of 7 negative controls (gray), and those with signal more than 3 standard deviations (SD) above the mean of these negative controls (dashed line) were considered to be detected above background (solid circles). Background-subtracted probeset reactivity profiles were compared with the reference panel (Supplementary Figure S4), and the closest match was taken as predicted species identity (correct predictions shown in red, incorrect in black). Limits of detection and distinction are tabulated for each species in **(f)**.

### Clinical blood cultures positive for yeast are correctly identified using the Phirst-ID assay directly from culture media

The Phirst-ID assay works directly from clinical samples, as it requires no enzymology, only hybridization, and because the rRNA sequences that form the basis of species identification are invariant to physiological perturbations (18). Given the high clinical importance of candidemia, we piloted *Candida* Phirst-ID on 62 consecutive positive blood culture bottles at two local hospitals whose Gram stain showed yeast. Briefly, broth from each blood culture bottle was lysed and run directly on the NanoString (see Methods), and the reactivity pattern of each sample was compared with that of our reference panel via Pearson correlations, exactly as for the set of 41 test strains grown in laboratory culture. In 60 of the 62 samples, the clinical microbiology laboratory identified a *Candida* species via their standard workflow, of which 59 were among the species in our reference panel, and one was not (*C. melibiosica*); the two non-*Candida* species included one *Saprochaete clavata* and one *Rhodotorula mucilaginosa* (**Supplementary Table S4**). For 58 of the 59 *Candida* species included in our reference panel, Phirst-ID matched the species identified by the clinical microbiology laboratory (**Figure 4**; 98.3% concordance, 95% confidence interval 92.3-99.8%). The discordant sample (BFx121) ended up being a mixed culture, though this was not recognized by the clinical microbiology laboratory. Of the remaining three yeast that were not in the reference panel, none showed substantial cross-reactivity with any of the species-specific probes (relative signal from the maximal species-specific probe <0.5% of the maximal genus-specific probe, compared with >9% for all tested species in the panel). *C. melibiosica* exhibited a pattern of reactivity to both pan-*Candida* probes that was similar to the other 59 *Candida* species, whereas *S. clavata* showed a unique hybridization pattern, with an inverted signal intensity between the two probes compared with any of the 60 *Candida*, and *R. mucilaginosa* was only recognized by the 28S pan-*Candida* probe, not the 18S. Thus, it was clear from inspection of the binding heatmap that those three specimens did not match any species in our reference panel, although the pan-*Candida* probes still indicated the presence of a yeast species (even for two non-*Candida* yeasts, albeit with atypical reactivity patterns).

**Figure 4.**
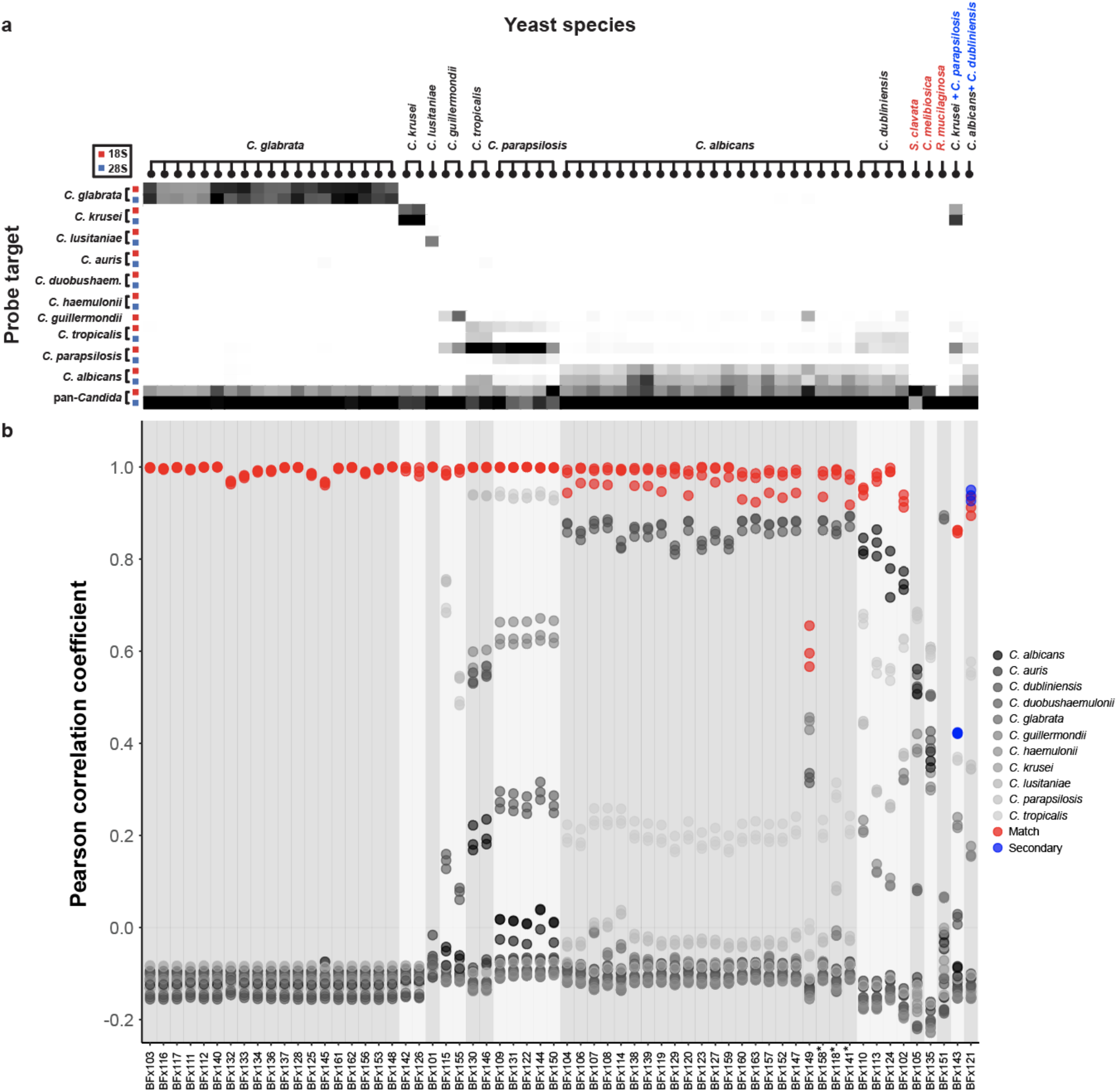
Clinical blood cultures identified using *Candida* Phirst-ID. Phirst-ID signal from 62 consecutive clinical blood culture specimens that showed yeast on Gram stain, shown as **(a)** heatmaps of normalized, background-subtracted signal, scaled to the maximum signal in each sample and **(b)** Pearson correlation coefficients of probeset reactivity profiles compared with the reference panel. Shading of data points indicate the comparison species from the reference panel; species matches are shown in red. Samples are labeled above and ordered by species identity assigned by the clinical microbiology laboratory, except for two samples at right that showed mixed binding intensities by Phirst-ID; for these, the correlation coefficient for the second species identified is shown in blue. Red species labels indicate off-panel organisms. Asterisks below indicate samples that also grew bacteria.

### Phirst-ID assay detects mixed blood culture specimens

We noted by visual inspection of binding heatmaps that two samples (BFx121 and BFx143), including the one with discordant identification, appeared to have additional signals beyond those expected from the best-matched species (**Figure 4; Supplementary Figure S5a**). Since both appeared to be explained by additive signals from two different *Candida* species, we hypothesized that these might represent mixed cultures, though only one species was reported by the clinical microbiology laboratories. Remarkably, both patients from whom these samples were obtained grew another *Candida* species from blood culture at a different time in the same hospital admission; our hybridization data suggested that this additional species may have been present at a low level in the sample we tested in each case. For the one discordant sample (BFx121), Phirst-ID identified *C. dubliniensis*, while the clinical microbiology lab identified *C. albicans* from the blood culture bottle we obtained. However, this patient also grew *C. dubliniensis* from other bottles drawn simultaneously, though we were only given one of these bottles to process, per our protocol. This patient had tricuspid valve endocarditis, and a vegetation from that valve grew both *C. albicans* and *C. dubliniensis*; our Phirst-ID assay showed signals from both species in this one blood culture bottle. In one other sample that appeared to clearly have mixed signal despite a concordant identification (BFx143), the patient grew *C. krusei* from the specimen in question, but had grown *C. parapsilosis* from a specimen drawn 8 days earlier. In both cases, the samples in question appeared to have Phirst-ID signals that corresponded to a linear combination of the species identified in the clinical microbiology laboratory (*C. albicans* for BFx121 and *C. krusei* from BFx143), as well as the other species grown by that patient at another time in their hospitalization (*C. dubliniensis* for BFx121 and *C. parapsilosis* for BFx143).

To test our hypothesis that these two specimens might represent mixed cultures, we streaked the blood culture broth from each specimen onto ChromAgar *Candida* plates (**Supplementary Figure 5b**). In one case (BFx143), two distinct colony morphologies were observed. We ran each morphotype on the Phirst-ID assay and found that the pink morphotype matched *C. krusei*, the species assigned by both the bulk Phirst-ID assay and the clinical microbiology lab, whereas the white morphotype better matched *C. parapsilosis*, which corresponded to the additional binding density seen visually in the bulk Phirst-ID assay, and the species grown at different times during that patient’s hospitalization (**Supplementary Figure 5c**). In the other case (BFx121), only one morphotype was observed, but both *C. albicans* and *C. dubliniensis* form light green colonies on ChromAgar *Candida* plates that are often indistinguishable (27). We therefore sampled 8 colonies at random and ran them on the NanoString, and found that 6 showed PSRPs consistent with *C. albicans* while 2 showed PSRPs consistent with *C. dubliniensis*, confirming the suspected mixture of these two species in the blood culture broth. For each colony type from both BFx121 and BFx143, we also sequenced the ITS region of each representative colony by Sanger sequencing, which confirmed the Phirst-ID results, showing *C. albicans* and *C. dubliniensis* from BFx121 and *C. krusei* and *C. parapsilosis* from BFx143. A third culture, BFx149, also had an unexpectedly high reactivity to the *C. guilliermondii* probe despite otherwise best matching *C. albicans*, albeit with a lower Pearson correlation than the rest; it was identified in the clinical microbiology laboratory as *C. albicans*. On restreaking this culture, only one morphotype was seen, and nine randomly selected colonies all matched *C. albicans* by Phirst-ID and Sanger sequencing of the ITS region; the *C. guilliermondii* probe reactivity seen from blood culture broth was not seen in any of these individual colonies.

In addition to the 62 clinical specimens, one post-mortem blood culture (BFx154.PM) was received by the BWH clinical microbiology lab during our period of sample collection that showed both yeast and bacteria on Gram stain. It also demonstrated what appeared to be a mixture of signals from *C. krusei* and *C. glabrata* by Phirst-ID (**Supplementary Figure S5a**); the clinical microbiology laboratory identified only *C. krusei*. We streaked this on ChromAgar *Candida* as well, again revealing two morphotypes (**Supplementary Figure S5b**), one of which resembled *C. krusei* (pink, fuzzy colonies) and the other *C. glabrata* (pink, smooth). These species identities were confirmed by *Candida* Phirst-ID on the independent isolates (**Supplementary Figure S5c**), again demonstrating that Phirst-ID can detect mixed *Candida* populations in blood culture bottles.

Finally, three clinical blood cultures (BFx118, 141, and 158), and the post-mortem sample (BFx154.PM), grew both yeast and bacteria in the same bottle (indicated with asterisks in **Figure 4**). Conceptually, the Phirst-ID approach can detect each of these microbes, and the sequences of bacterial and fungal rRNA diverge enough that cross-reactivity or assay interference would not be expected. Consistent with this, the *Candida* species in each of these samples was identified accurately. Since we previously designed a bacterial Phirst-ID probeset (18), we also tested an aliquot of each of these clinical specimens using this bacterial probeset, as well as two specimens that were not reported to grow any bacteria, only yeast. In each of the specimens from which bacterial growth was reported, we found reactivity profiles from the bacterial Phirst-ID probeset consistent with the species reported from the clinical microbiology laboratory (**Supplementary Figure S6**), including *Staphylococcus epidermidis* (BFx118 and 158) and *Enterococcus faecium* (BFx141) from the clinical specimens, and a combination of a strong signal from *Enterococcus faecalis* and faint signal from *Klebsiella pneumoniae* from the post-mortem specimen (BFx154.PM). No bacterial Phirst-ID signals were found from the two specimens for which no bacterial growth was reported (BFx124 and BFx150).

## Discussion

Candidemia is one of the top 10 causes of bloodstream infection in hospitalized patients, with high attendant mortality that rises with delays in treatment initiation (7, 9, 10, 28, 29). Current clinical diagnostic methods for invasive fungal infections require considerable expertise, infrastructure, and time. Even with the advent of mass spectrometry-based pathogen identification, delays in clinical diagnosis often result from slow subculture steps required for optimal accuracy (5, 12). We recently developed Phirst-ID, a novel approach for amplification-free, multiplexed fluorescence-based rRNA hybridization to identify bacteria directly from clinical specimens (18). Here, we extend this Phirst-ID approach, designing a 21-plex probeset that can rapidly and accurately identify 11 common pathogenic *Candida* species from both isolated colonies (33/33 on a reference panel, 33/33 in an independent validation set) and positive clinical blood culture broths (concordant identification of 58/59 *Candida* species included in the reference panel, where the only discordant sample was a mixed specimen that included the species identified by Phirst-ID; no mis-identifications of 3 additional yeast species that were not in the reference panel).

Phirst-ID is straightforward to perform, requiring <30 minutes of hands-on time and <8 hours total assay time on our current instrument, the NanoString Sprint; recent advances in instrumentation and optics can accelerate the workflow to <4 hours on a pilot instrument (30). By comparison, identification of the yeast in these samples by standard clinical microbiology workflows took a median of 28 hours (range 14 - 36) from the time the blood cultures signaled positive. The lack of any enzymatic step means that Phirst-ID works directly on crude lysate without nucleic acid purification, reducing assay complexity. Further, since experimental conditions need only to permit base pairing, chemical RNase inhibitors (here, ß-mercaptoethanol) can be used without interfering with the assay, enhancing robustness compared with other RNA-based techniques. The rich phylogenetic information encoded in rRNA sequences, along with their extreme conservation within species, have long made them an ideal target for species identification (31, 32). And since Phirst-ID is based only on rRNA sequence, not relative expression, it should be invariant to physiological conditions, obviating the need for subculture. Further, as the most abundant nucleic acid (many thousands of copies per cell), rRNA detection enables highly sensitive pathogen detection even without an amplification step: we were able to identify species from fewer than 30 cell equivalents per assay, and to detect the presence of a *Candida* species from roughly 1 cell equivalent.

The detection of two mixed fungal samples among 62 consecutive blood cultures with yeast, neither of which were recognized to be mixed by standard workflow in the clinical microbiology laboratory, was unexpected. We saw no evidence of mixed signals in 66 samples prepared from laboratory culture, so this is not likely to represent an assay artifact; further, we were able to isolate individual colonies of each of the species that contributed to the mixed binding signals by streaking the blood culture broth onto ChromAgar *Candida* plates, supporting the findings of the primary assay. Also, each patient grew the second species identified by Phirst-ID from blood at other times in their hospital course, supporting the clinical significance of our findings. The ability of Phirst-ID to identify such mixed species, a challenge for other assays that work directly from blood culture without a subculture step (33, 34), thus may be an unanticipated strength of this approach, and the frequency of mixed *Candida* species in blood cultures should be further explored.

This approach has several limitations. First, while it proved accurate at identifying *Candida* species that were present in the reference panel, it will be able to discriminate but not identify yeast species which it has not previously encountered, as evidenced by the three blood cultures growing *C. melibiosica* and two non-*Candida* species (*S. clavata* and *R. mucilaginosa*). In order to more broadly identify other yeast in blood, the probeset design will need to be extended to other species and to include additional genus level classification probes, and the reference panel will need to be expanded as well. Iterative expansion is entirely feasible, and our prior results with a 180-plex rRNA-targeted bacterial probeset demonstrates that the assay can tolerate far more complexity if needed (18). Second, although this assay was able to detect several instances of mixed cultures, this required visual inspection of the reactivity pattern. Further work would be required to flag samples with anomalous patterns (e.g., lower-than-expected correlation coefficients) for manual review, or ultimately to automate recognition of linear combinations of reactivity patterns from multiple species. Third, despite a limit of detection around 1 cfu, as currently configured, Phirst-ID still requires an initial culture step from blood, due to limitations in sample preparation and pathogen concentration. Finally, this assay remains research-use only for now; additional work will be required to streamline, standardize, and ideally automate sample processing, overall workflow, analysis, and interpretation prior to clinical implementation. Still, the accuracy achieved from this pilot assay on clinical blood cultures was promising.

Given demonstrated past success on identification of bacteria (18), viruses, and parasites (35), and these results in *Candida*, this approach holds the potential for pan-microbial identification in the future. Indeed, in each of four blood cultures that grew both *Candida* and bacteria, separate Phirst-ID probesets were able to detect each pathogen class. In principle, Phirst-ID probesets for bacteria, fungi, and other pathogens could be combined in a single diagnostic assay. Thus, with further probeset design, optimization, and combination, hybridization-based, rRNA-targeted identification strategies show promise for rapid, accurate, and highly sensitive detection of pathogens directly from clinical samples, as well as in agricultural, environmental, or biodefense applications.

## Acknowledgments

We thank Richard Bennett and the CDC ARBank for sharing strains used in the reference panel. This work was supported in part by the National Institute of Allergy and Infectious Diseases of the National Institutes of Health (award numbers 1R01AI153405 to R.P.B. and T32 AI007061 to M.E.M.) and the Massachusetts General Hospital Transformative Scholars Program (R.P.B.). The content is solely the responsibility of the authors and does not necessarily represent the official views of the National Institutes of Health. The funders had no role in study design, data collection and interpretation, or the decision to submit the work for publication. R. Boykin is an employee at NanoString, Inc., the company that manufactures the RNA detection platform used in this manuscript. R.P.B. is a co-inventor on subject matter in PCT/US2014/027158 and PCT/US2014/068835, filed by the Broad Institute directed to rRNA hybridization for organism identification, and an accelerated method for rRNA hybridization, respectively, as described in this manuscript. NanoString, Inc. has licensed the intellectual property for rRNA hybridization-based organism identification from the Broad Institute.

## SUPPLEMENTARY TABLES

**Supplementary Table S1.** 18S and 28S sequences from (a) ingroup and (b) outgroup species used in probeset design

**Supplementary Table S2.** Probe sequences

**Supplementary Table S3.** Strain list from (a) reference and (b) validation strain panels

**Supplementary Table S4.** Clinical samples

## SUPPLEMENTARY FIGURE LEGENDS

**Supplementary Figure S1.**
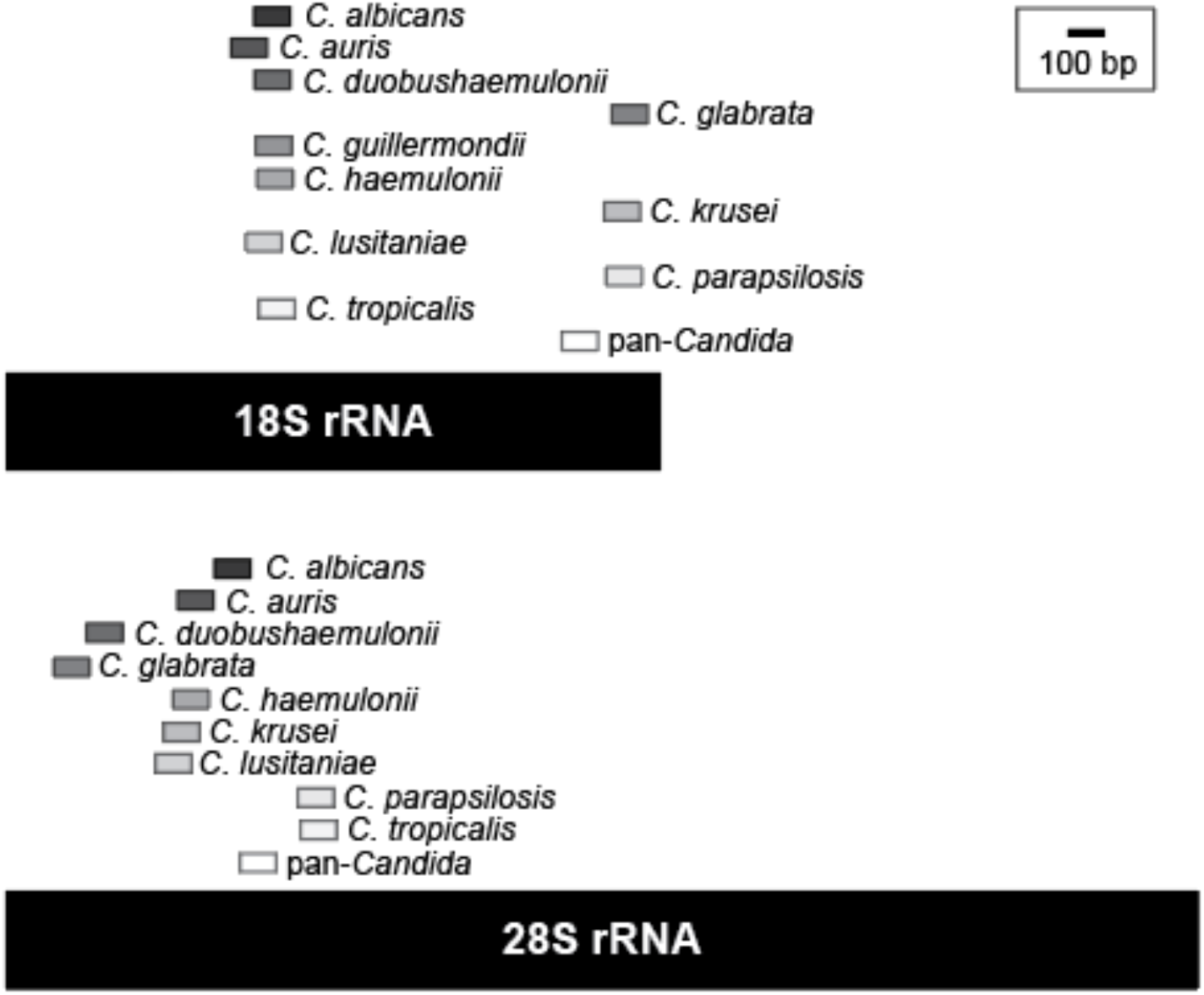
Target regions of 21 *Candida* Phirst-ID probes on 18S and 28S rRNA subunits. Each of the 21 probe pairs in the *Candida* Phirst-ID probeset are mapped onto their complementary regions in schematic representations of the 18S or 28S rRNA subunits.

**Supplementary Figure S2.**
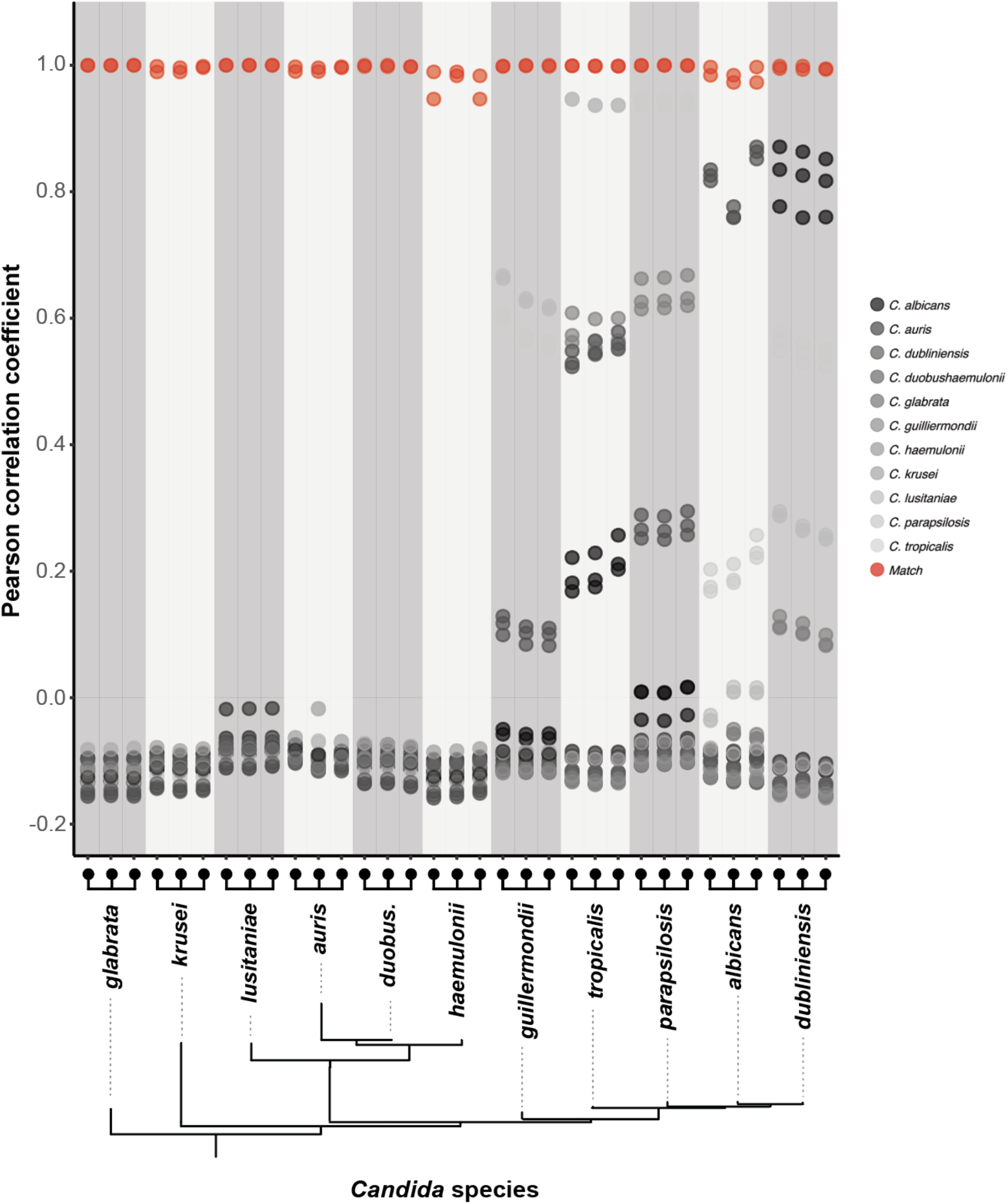
Non-self Pearson correlation coefficients of *Candida* Phirst-ID profiles identify species in a reference panel. Pearson correlation coefficients of probeset reactivity profiles of species-specific probes from 3 isolates of each of 11 species in a reference panel, against each other panel member, are plotted. Shading of data points indicate the comparison species; species matches are shown in red.

**Supplementary Figure S3.**
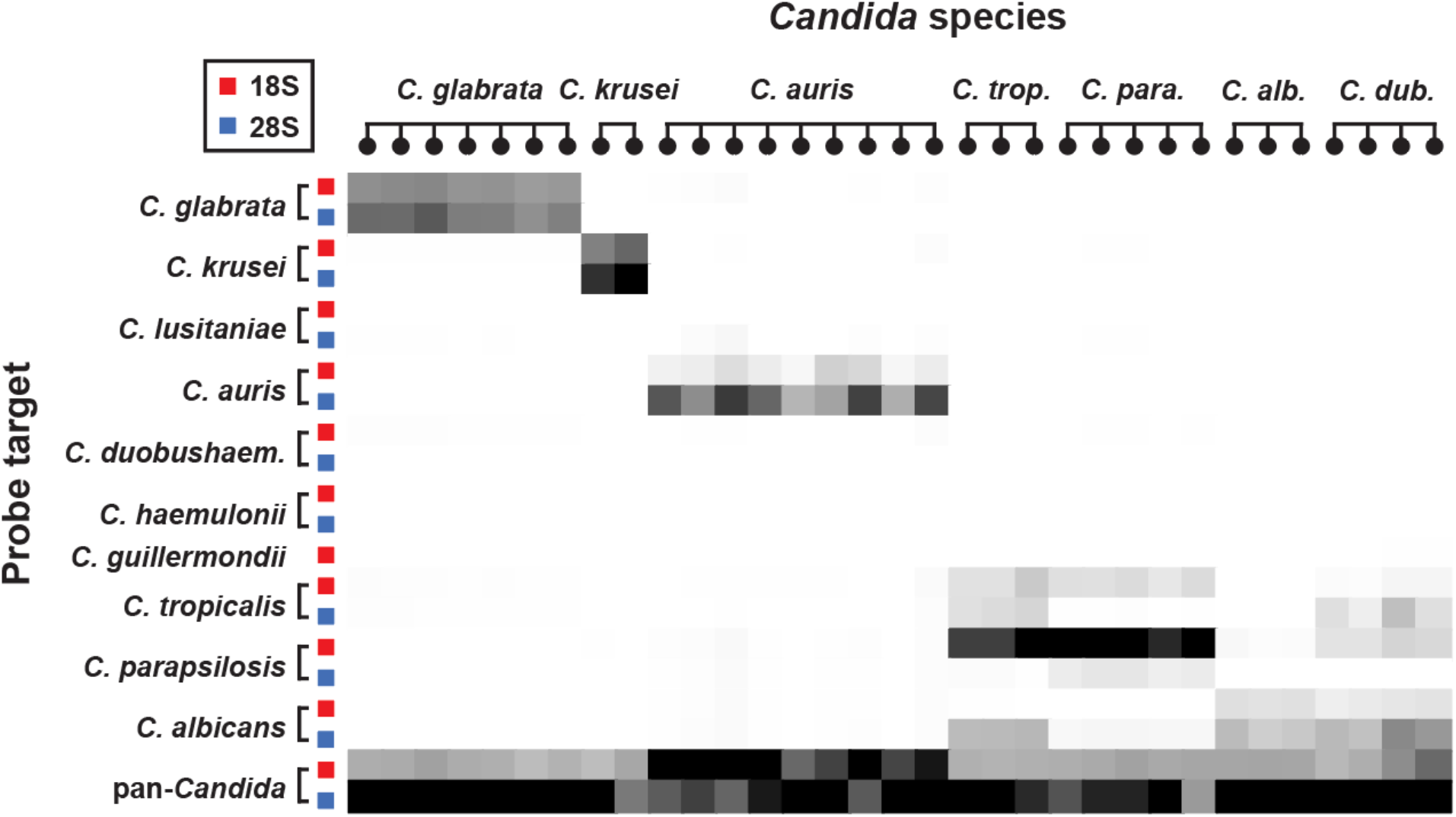
Independent validation of *Candida* Phirst-ID probeset. Heatmap of normalized, background-subtracted binding intensities show Phirst-ID probeset reactivity profiles of 33 isolates from 7 *Candida* species grown in laboratory culture, independent of the reference panel in Figure 1a. Heatmap intensity is normalized within each sample to the maximum signal for that sample.

**Supplementary Figure S4.**
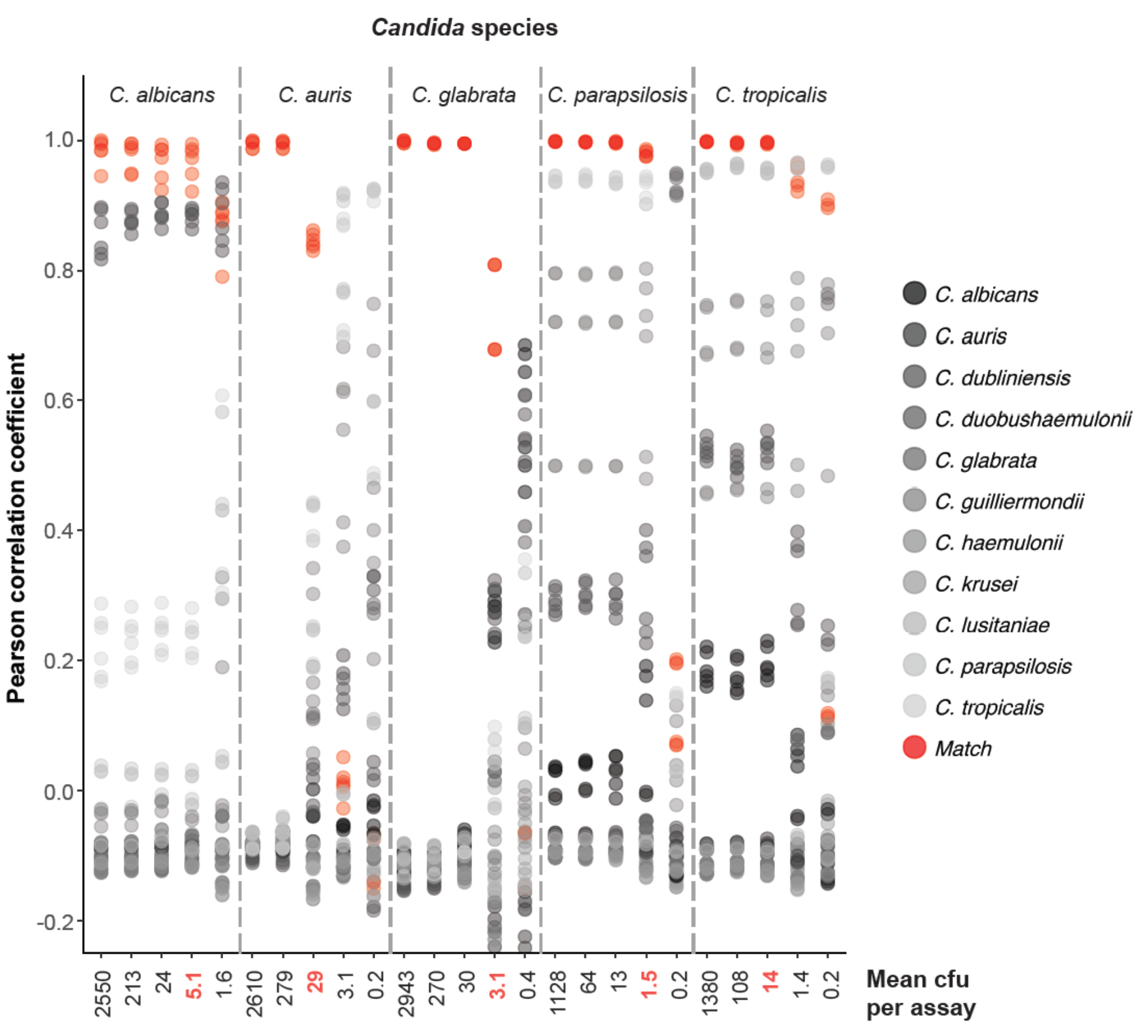
Phirst-ID limits of distinction for 5 *Candida* species. Pearson correlation coefficients of Phirst-ID probeset reactivity profiles from serial dilutions of each indicated *Candida* species (upper horizontal axis label) versus the reference panel, with cfu for each sample as indicated (lower horizontal axis label). Shading of data points indicate the comparison species from the reference panel; species matches are shown in red. The lowest cfu per assay at which the highest Pearson correlation coefficient corresponded to the correct species is indicated in red as the measured limit of distinction for that species.

**Supplementary Figure S5.**
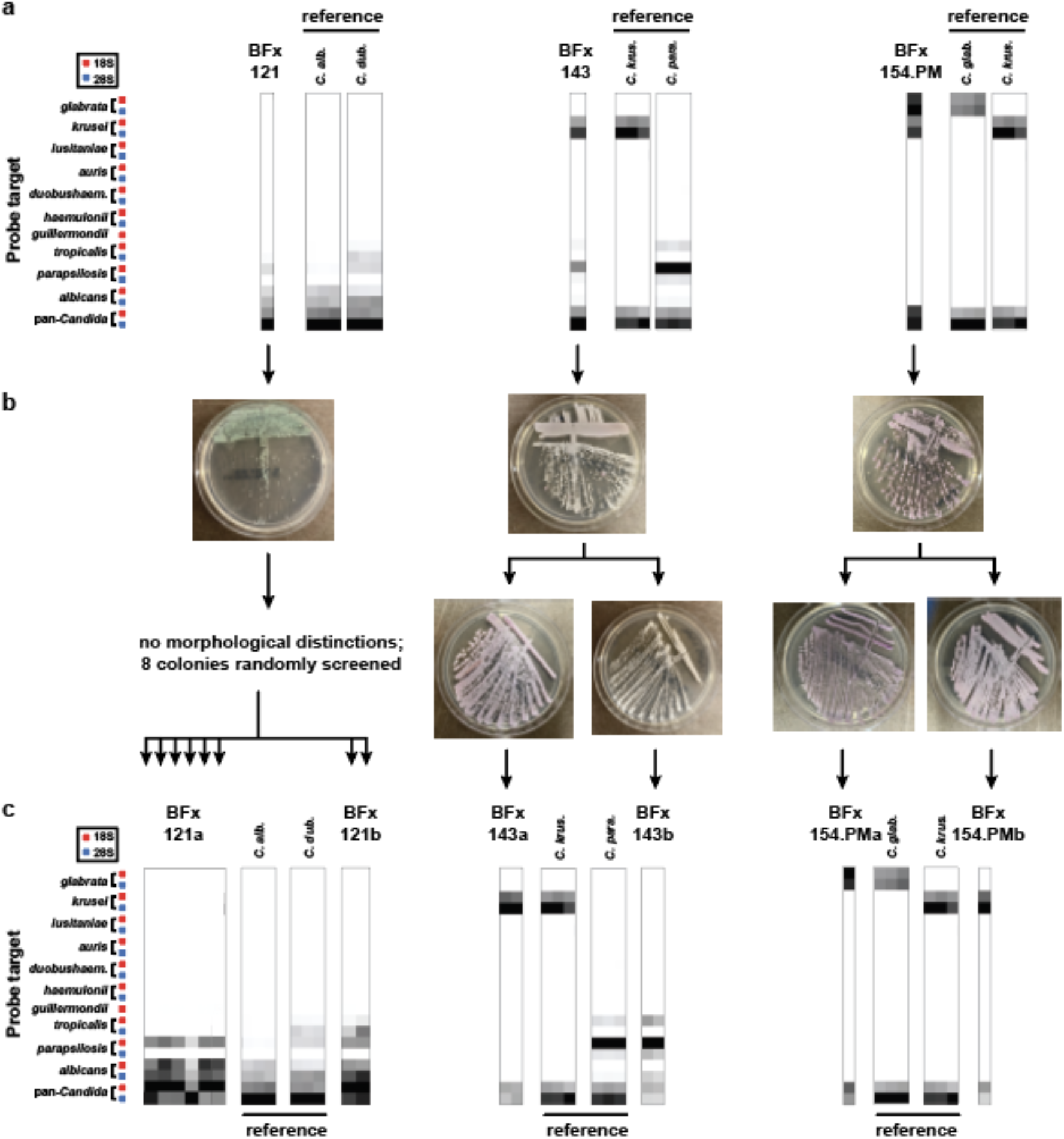
*Candida* Phirst-ID revealed mixed blood cultures that were missed by standard clinical microbiology workflows. **(a)** Phirst-ID probeset reactivity profiles (PSRPs) for two clinical blood cultures (BFx121 and BFx143) and one post-mortem blood culture (BFx154.PM) appeared to be a linear combination of two species. PSRPs from three reference panel isolates from the two species that appeared to be represented are shown at right. **(b)** Blood culture broth from each of these three samples was streaked onto ChromAgar *Candida* plates, in two cases revealing two distinct culture morphologies (top row) that could be restreaked to homogeneity from single colonies of each morphotype (bottom row). **(c)** Phirst-ID PSRPs from individual colonies unambiguously matched only one reference species, confirming that each original blood culture broth represented a mixture of two species. PSRPs from three reference panel isolates of each matching species are again provided for comparison.

**Supplementary Figure S6.**
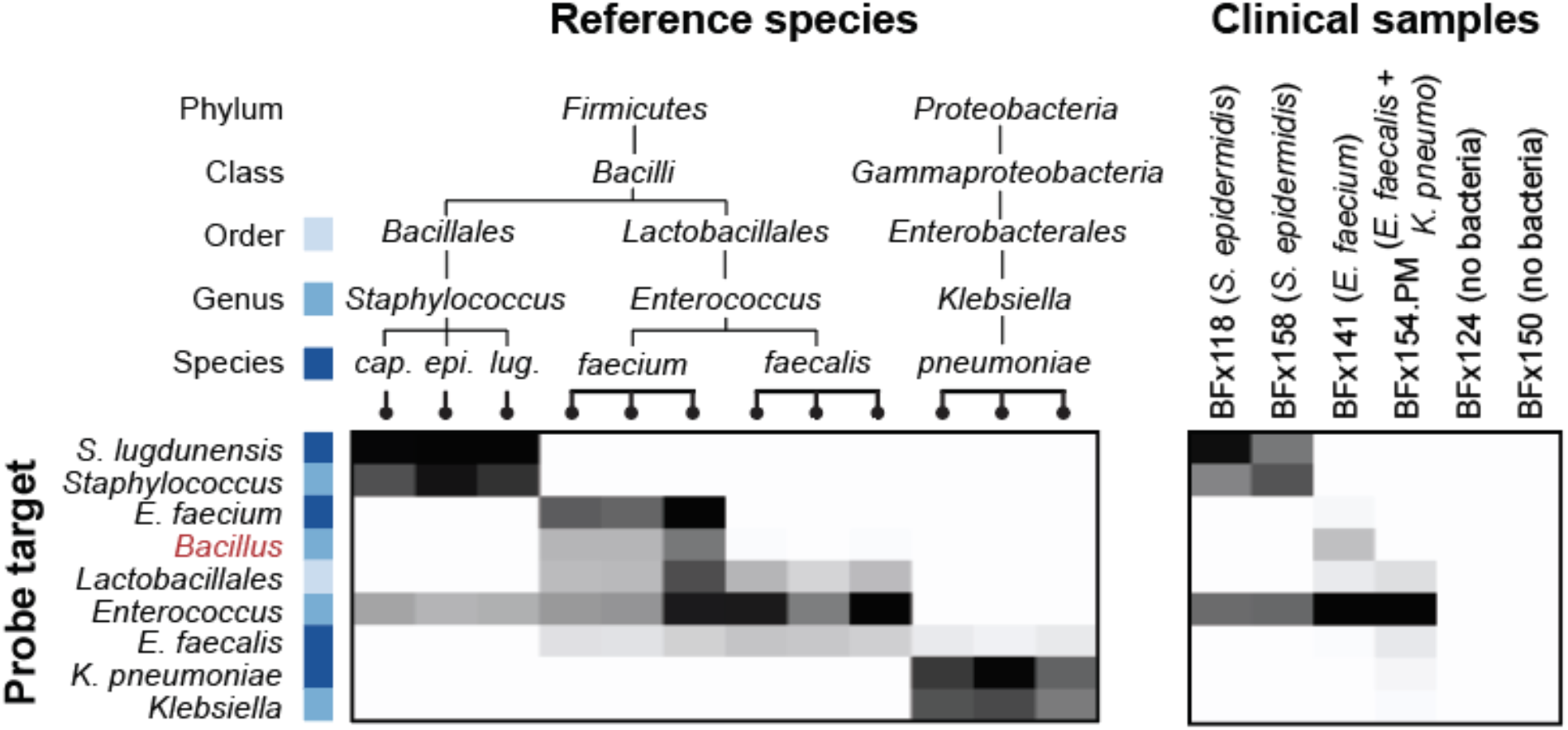
Bacterial Phirst-ID recognizes bacteria from mixed fungal and bacterial blood cultures. Left panel: bacterial Phirst-ID data from selected probes for reference strains are shown at left (from Bhattacharyya et al, *Sci Rep* 2019, reference 18), with phylogenetic hierarchy of probe target encoded in color. Red probe = off-target probe designed against *Bacillus* genus that fortuitously distinguished *E. faecium* from *E. faecalis* due to unanticipated cross-reactivity. Right panel: data from the same selected bacterial Phirst-ID probes for 6 clinical or post-mortem blood cultures that grew yeast, 4 of which also grew bacteria, and 2 of which did not. Bacterial identifications for each sample from the clinical microbiology laboratory are shown in parentheses. Heatmaps display normalized, background-subtracted read intensities for each probe, scaled to the maximum signal in each sample. (*cap.* = *capitis*, *epi.* = *epidermidis*, *lug.* = *lugdunensis*)

